# Single-cell RNA sequencing reveals a role for reactive oxygen species and peroxiredoxins in fatty-acid-induced rat ß-cell proliferation

**DOI:** 10.1101/2021.12.22.473904

**Authors:** Alexis Vivoli, Julien Ghislain, Ali Filali-Mouhim, Zuraya Elisa Angeles, Anne-Laure Castell, Robert Sladek, Vincent Poitout

**Author notes:** Corresponding author and lead contact; @VPoitout.

## Abstract

The functional mass of insulin-secreting pancreatic β cells expands to maintain glucose homeostasis in the face of nutrient excess, in part via replication of existing β cells. To decipher the underlying molecular mechanisms, we assessed β-cell proliferation in isolated rat islets exposed to glucose and oleate or palmitate for 48 h and analyzed the transcriptional response by single-cell RNA sequencing. Unsupervised clustering of pooled β cells identified subpopulations, including proliferating β cells. β-cell proliferation increased in response to oleate but not palmitate. Both fatty acids enhanced the expression of genes involved energy metabolism and mitochondrial activity. Comparison of proliferating vs. non-proliferating β cells and pseudotime ordering suggested the involvement of reactive oxygen species (ROS) and peroxiredoxin signaling. Accordingly, the antioxidant N-acetyl cysteine and the peroxiredoxin inhibitor Conoidin A both blocked oleate-induced β-cell proliferation. Our data reveal a key role for ROS signaling in β-cell proliferation in response to nutrients.

## INTRODUCTION

Both type 1 and type 2 diabetes are characterized by a loss of functional β-cells (Chen et al., 2017, Weir et al., 2020), and increasing β-cell mass is considered a promising therapeutic approach (Zhou and Melton, 2018). In type 2 diabetes, a commonly accepted view is that insulin resistance drives a compensatory increase in both insulin secretion from individual β cells and β-cell numbers (Esser et al., 2020), in part via replication of existing β cells (Dor et al., 2004). This view is supported by the observations that insulin (Okada et al., 2007) and factors secreted from peripheral tissues in response to insulin resistance, such as serpinB1 (El Ouaamari et al., 2016), promote β-cell proliferation. However, the notion that insulin resistance is the cause of compensatory β-cell mass expansion has been challenged (Johnson, 2021). In humans, insulin hypersecretion can be observed prior to insulin resistance in obesity (van Vliet et al., 2020), and in mice, short-term exposure to high-fat diet leads to enhanced β-cell proliferation before an apparent increase in insulin resistance (Mosser et al., 2015, Stamateris et al., 2013). This suggests a direct effect of nutrients to promote β-cell replication. Accordingly, short-term glucose infusion in mice leads to a significant increase in β-cell proliferation (Alonso et al., 2007), and we previously showed that a combination of glucose and fatty-acids (FA) increases β-cell proliferation and mass in infused rats as well as in rat and human islets ex vivo (Fontés et al., 2010, Moullé et al., 2017).

As islets contain several cell types and β cells are functionally and transcriptionally heterogeneous (Wang and Kaestner, 2019), we utilized single-cell RNA sequencing (scRNA-seq) to decipher the molecular mechanisms underlying FA-induced β-cell proliferation. Specifically, the aims of this study were 1) to ascertain the differential effects of individual FA on β-cell replication; and 2) to characterize the transcriptome of proliferating β-cells in response to FA at single-cell resolution.

## RESULTS

### scRNA-seq identifies a subpopulation of proliferative β cells

Isolated rat islets were cultured for 48 h in the presence of 16.7 mM glucose (Vehicle, n=4) with or without 0.5 mM of the saturated FA palmitate (C16:0, n=3) or the monounsaturated FA oleate (C18:1, n=4), the 2 most abundant circulating FA in humans (Abdelmagid et al., 2015) (Figure 1A). Following scRNA-seq, unbiased graph-based clustering of 52,570 pooled cells from all 3 culture conditions identified the 4 main endocrine cell types based on expression of insulin (*Ins2*, n = 39,910), glucagon (*Gcg*, n = 7,821), somatostatin (*Sst*, n = 1,419) and pancreatic polypeptide (*Ppy*, n = 1,157), corresponding to β, δ and PP cells, respectively, as well as non-endocrine cells (n = 2,263) (Figure 1B, Suppl. Figure 1A). Endocrine cell types clustered together irrespective of the treatment condition. Refined clustering of the *Ins2*-positive cells identified 5 β subpopulations: Beta 1 (n = 28,411), Beta 2 (n = 638), Beta 3 (n = 5,811), Beta 4 (n = 3,596) and Beta 5 (n = 1,454) (Figure 1C). Except for cluster Beta 4, where only 10 genes (including *Ins1* and *Ins2*) involved in hormone activity were enriched (Suppl. Figure 1B, Suppl. Tables 2 & 3), the other clusters displayed a large number of uniquely enriched genes [log_2_ fold change ≥ 0.25: 227 for Beta 1, 1,063 for Beta 2, 611 for Beta 3 and 662 for Beta 5 (Suppl. Table 2)]. Differentially expressed genes (DEGs) and pathway enrichment analysis from the top 50 cluster markers for the Beta 1 subpopulation suggest a role in hormone secretion (e.g., *Ptprn2*, *Vamp2*) (Suppl. Figure 1B, Suppl. Tables 2 & 3). Genes involved in cell division processes were highly enriched in the Beta 2 subpopulation, as shown by expression of the proliferation markers *Mki67*, *Cdk1* and *Pcna* (Figure 1D, Suppl. Figure 1B, Suppl. Tables 2 & 3). The Beta 3 cluster was characterized by enriched expression of genes involved in the unfolded protein response (UPR) and endoplasmic reticulum (ER) stress including *Hspa1b*, *Ddit3* and *Tmbim6* (Suppl. Figure 1B, Suppl. Tables 2 & 3). Finally, the Beta 5 cluster was enriched in genes implicated in proinsulin biosynthesis and secretory vesicle formation as shown by the expression of *Scg2*, *Pcsk2*, *Chga* and *Cpe* (Suppl. Figure 1B, Suppl. Tables 2 & 3). Taken together, these results confirm the capture of heterogeneous β-cell clusters, including rare proliferative cells.

**Figure 1.**
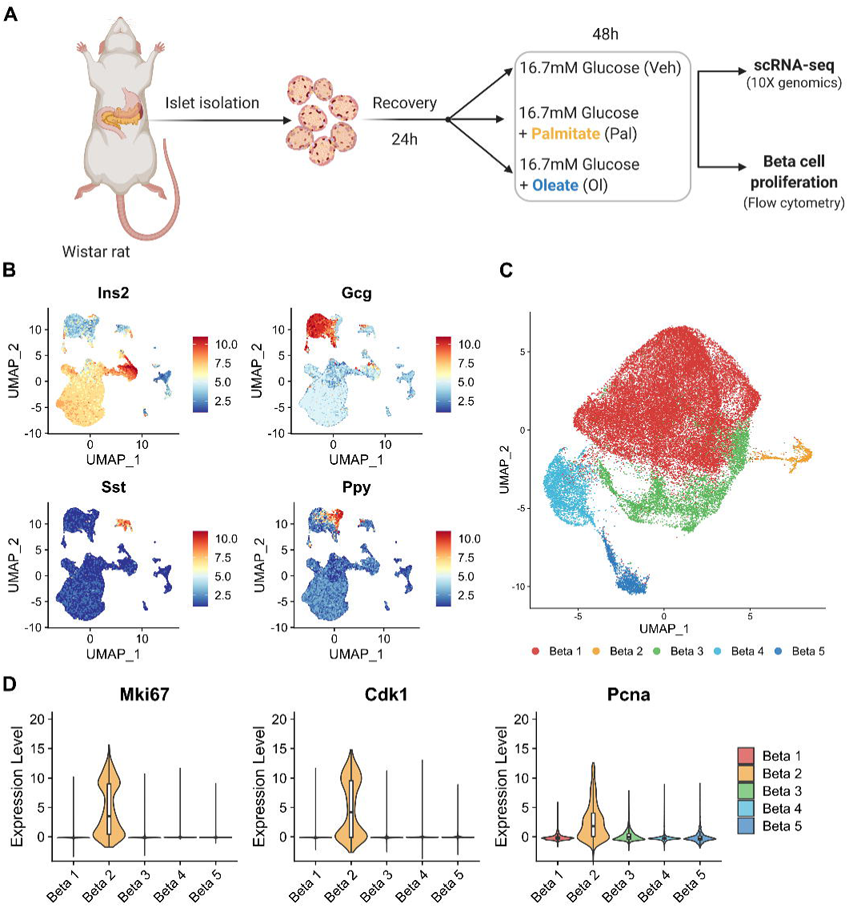
Single cell transcriptome analyses of rat islets. **(A)** Experimental design. Islets were isolated from 2-month-old male Wistar rats, allowed to recover for 24 h and cultured for 48 h with 16.7 mM glucose with vehicle (Veh), 0.5 mM palmitate (Pal) or oleate (Ol) and harvested for scRNA-seq or flow cytometry. Image created at Biorender.com. **(B)** UMAP plots of pooled (Veh, Pal and Ol conditions) endocrine cells showing expression of insulin (*Ins2*), glucagon (*Gcg*), somatostatin (*Sst*), and pancreatic polypeptide (*Ppy*). **(C)** UMAP plot of pooled (Veh, Pal and Ol conditions) β cells showing the five β-cell clusters (Beta 1 to Beta 5). **(D)** Violin plots showing the expression of proliferative markers in the β-cell clusters. Boxplot in violin interior shows median, quartile and whisker values.

### Oleate and palmitate differentially impact the **β**-cell transcriptome

We compared the transcriptomes of individual β-cell clusters between oleate- or palmitate-and vehicle-treated islets and performed pathway enrichment analysis (Figure 2).

**Figure 2.**
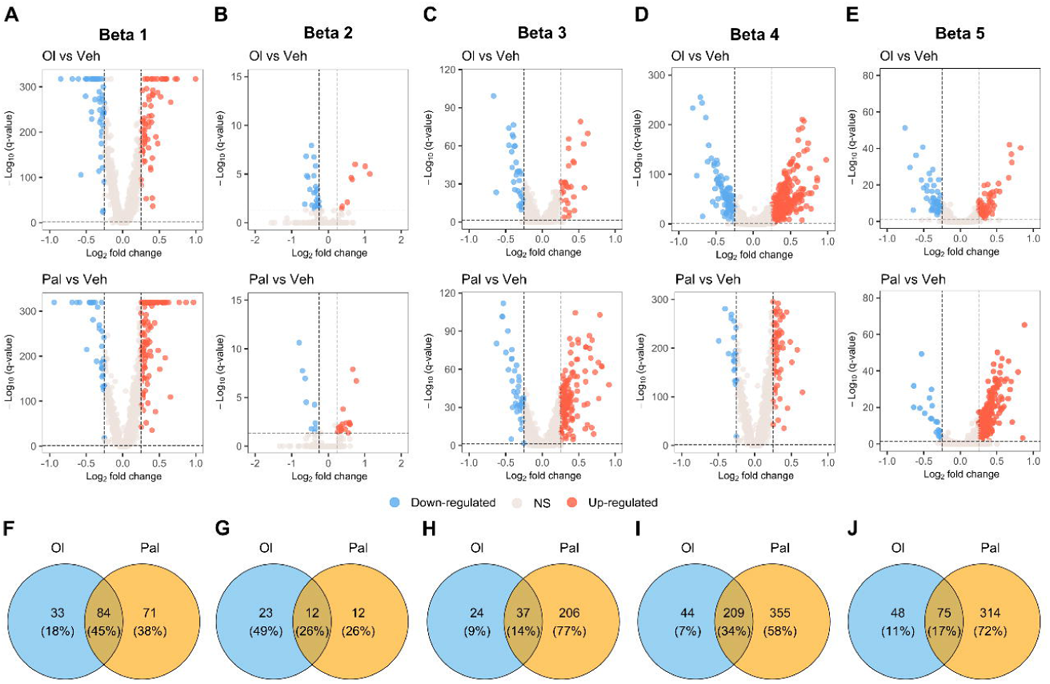
Differentially expressed genes in β-cell clusters. **(A-E)** Volcano plots showing the Log_2_ fold change of gene expression and the statistical significance (-log_10_(q-value) <0.05) in the Beta 1-Beta 5 from Ol- or Pal- vs Veh treated islets. Upregulated (red) and downregulated (blue) genes are shown. **(F-J)** Venn diagrams shows the number of shared or unique regulated genes [log_2_ (fold change) > 0.25, −log_10_(q-value) <0.05] in β-cells subpopulations.

For the Beta 1 cluster, oleate treatment resulted in 117 DEGs whereas palmitate treatment produced 155 DEGs. About 45% of the DEGs were similar between both conditions (Figures 2A & F). Both oleate and palmitate upregulated pathways controlling metabolic processes including glycolysis (e.g., *Aldoa*, *Gapdh*, *Pgk1* and *Eno1*) and oxidative phosphorylation (e.g. *Ndufa5, Cox5b* and *Atp5b*) (Figure 2A, Suppl. Tables 4 & 5).

In the proliferative Beta 2 cluster, comparison of oleate- or palmitate- to vehicle-treated islets revealed only 35 and 24 DEGs, respectively (Figures 2B & G). Except for downregulation of ribosomal protein subunits (e.g., *Rps7*, *Rps3a* and *Rps24*) and their associated pathways in oleate-treated islets, no major differences were observed between conditions (Figure 2B, Suppl. Tables 4 & 5). These results suggest that proliferative β cells have a similar transcriptomic profile in all culture conditions.

The Beta 3 cluster showed clear differences between palmitate and oleate treatments. Oleate-treated islets presented a total of 61 DEGs (Figure 2C & H). Pathway analysis showed decreased expression of β-cell differentiation and maturity markers (e.g., *Neurod1*, *Slc2a2* and *Pdx1*) and increased expression of anti-apoptotic response genes (e.g. *Tmbim6*) (Figure 2C, Suppl. Tables 4 and 5). Palmitate-treated islets displayed 243 DEGs (Figure 2C & H) including downregulated β-cell markers (e.g., *NeuroD1*, *Pdx1* and *Slc2a2*) (Figure 2C, Suppl. Tables 4 & 5) and genes involved in the regulation of insulin secretion (e.g. *Ptprn2, Gnas* and *Rgs16*) (Figure 2C, Suppl. Tables 4 & 5). In addition, UPR, ER stress, oxidative stress, apoptotic pathways (e.g., *Ddit3*, *Atf4, Herpud1*, *Sod2*) and translation processes (e.g., *Eif1ad*, *Rpl37, Rps28)* were upregulated in response to palmitate but not oleate (Figure 2C, Suppl. Tables 4 & 5). These data suggest that palmitate induces more severe dedifferentiation and dysfunction than oleate in the Beta 3 cluster.

In the Beta 4 cluster, oleate- and palmitate-treated islets displayed 253 and 564 DEGs, respectively (Figures 2D & I). No downregulated pathways were enriched in either treatment condition. Similar to the Beta 1 cluster, upregulated pathways following exposure to FA included cellular respiration (e.g., *Cox4i2*, *Ndufs3*, *Cycs* or *Atp5b*) and antioxidant response (e.g., *Sod2*, *Prdx1*) (Figure 2D, Suppl. Tables 4 & 5). However, transcripts involved in translation processes (e.g., *Eif1ad, Rpl37, Eif3d, Rpl41*) and ER stress-related pathways (e.g., *Ddit3*, *Ppp1r15a*, *Dnajb11*) were also enriched in palmitate-treated islets (Figure 2D, Suppl. Tables 4 & 5).

In the Beta 5 cluster, oleate exposure produced 123 DEGs (Figures 2E & J) including downregulation of β-cell differentiation and maturity markers (e.g., *Neurod1*, *Pdx1, Slc2a2* or *Nkx6-1)* and upregulation of pathways related to cellular respiration and antioxidant defense (e.g., *Cox6a1*, *Ndufs7*, *Sod2* or *Gpx1*) (Figure 2E, Suppl. Tables 4 & 5). On the other hand, palmitate-treated islets displayed 389 DEGs (Figures 2E & J). Similar to the Beta 3 and Beta 4 clusters, palmitate treatment increased expression of genes involved in oxidative phosphorylation (e.g., *Ndufs4*, *Cycs or Atp5d*), oxidative stress (e.g., *Sod2*, *Cycs*, *Gpx1*, *Prdx1*), translation processes (e.g., *Rpl41, Rps20*) and UPR/ER stress (e.g., *Ppia*, *Calr*, *Ddit3*, *Ppp1r15a*, *Dnajb11*) (Figure 2E, Suppl. Tables 4 & 5).

Taken together, these results reveal that while expression of genes involved in energy metabolism and mitochondrial activity pathways was upregulated in response to both FA, stress response pathways were only stimulated by palmitate. In addition, the absolute number of genes differentially regulated in response to palmitate was higher than in response to oleate. Finally, both FA decreased expression of β-cell identity genes in several clusters.

### Oleate stimulates β-cell proliferation

In the same batches of islets used for scRNA-seq, the proportion of proliferating β cells was determined by flow cytometry using the proliferation marker EdU. Oleate, but not palmitate, stimulated β-cell proliferation compared to vehicle (Figures 3A & B). Accordingly, analysis of the scRNA-seq data revealed a higher proportion of cells in the Beta 2 proliferating cluster in response to oleate (n=353) compared to palmitate (n=199) or vehicle (n=86) (Figure 3C).

**Figure 3.**
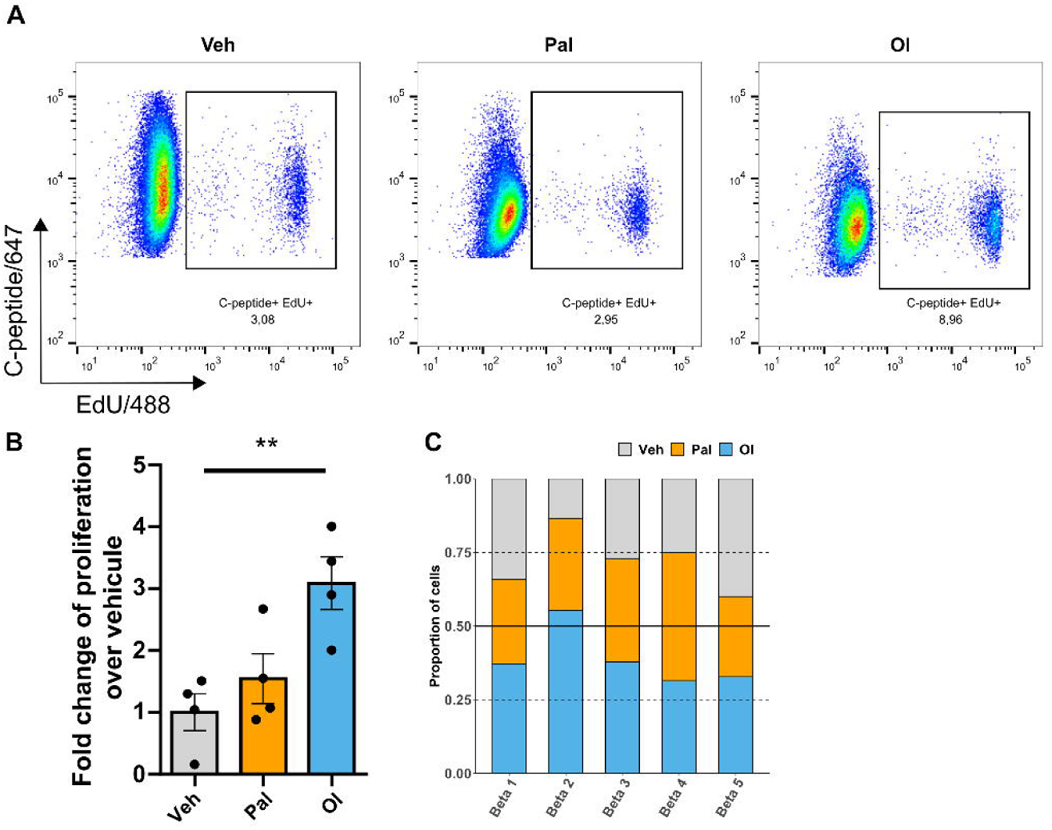
Analysis of β-cell proliferation. **(A, B)** Isolated rat islets were treated as describe in Figure 1A and β-cell proliferation assessed by flow cytometry following staining for EdU and C-peptide. **(A)** Representative dot plots showing the percentage of C-peptide^+^ EdU^+^ cells (boxed region) over total C-peptide^+^ cells in Veh, Pal and Ol treated islets. **(B)** Fold-change of proliferation over Veh condition. Data are expressed as means +/− SEM. **p<0.01 following one-way ANOVA with Dunnett’s multiple comparisons test. **(C)** Proportion of the five β-cell subpopulations identified by scRNA-seq in Veh, Pal and Ol-treated islets.

### Oxidative phosphorylation and oxidoreductase activity pathways are upregulated in proliferating **β** cells

scRNA-seq offers a unique opportunity to explore the transcriptome of proliferating β cells. Because no major transcriptional differences were detected in the proliferating Beta 2 cluster in response to oleate or palmitate compared to vehicle (Figure 2B), we compared the Beta 2 cluster to the non-proliferating Beta 1 and 3-5 clusters (Figure 4A, Suppl. Figure 2, Suppl. Table 6) irrespective of the treatment condition and performed pathway enrichment analysis. This approach enabled improved visualization of gene sets organized into networks. Comparison of Beta 2 with Beta 3 and Beta 5 showed downregulation of hormone and insulin response, glycosylation, cell secretion, UPR/ER-stress as well as lysosomal-related pathways (Suppl. Figures 2A & C). Comparison of Beta 2 to all non-proliferating clusters revealed that the majority of upregulated pathways were involved in protein catabolism and cell division-related processes, as expected (Figure 4A, Suppl. Figure 2). However, upregulation of glucose metabolism (Beta 2 vs. Beta 1, Figure 4A), ATP synthesis, oxidative phosphorylation, and oxidoreductase activity (Beta 2 vs. Beta 1 and 3-5, Figure 4A, Suppl. Figure 2) was also observed. Among these pathways, the main upregulated genes included the subunits of the complexes of the electron transport chain (e.g., *Ndufa12, Cyc1* and *Atp5g2*), involved in oxidative phosphorylation and ATP synthesis, and the antioxidant family members thioredoxin (e.g., *Txn1* and *Txnl1*) and peroxiredoxin (*Prdx1, Prdx2* and *Prdx4*), involved in oxidoreductase activity (Figure 4B). These findings highlight the importance of mitochondrial metabolism in the control of β-cell proliferation and suggest a role for reactive oxygen species (ROS) and antioxidant signaling.

**Figure 4.**
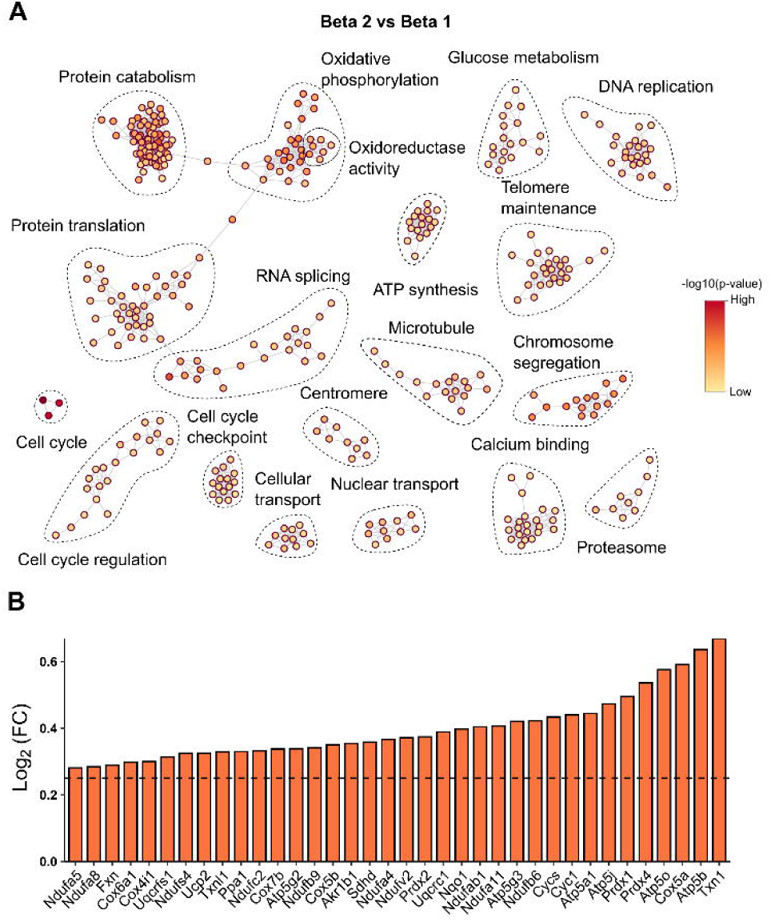
Comparison of proliferative to non-proliferative β cells. **(A)** Pathway enrichment map of pooled (Veh, Pa and Ol conditions) Beta 2 (proliferative) versus Beta 1 (non-proliferative) subpopulations. Nodes represent pathways. Gradient represents lower or higher significance for the up-regulated pathways. **(B)** Fold-change of expression of genes involved in oxidative phosphorylation and oxidoreductase activity related pathways in the Beta 2 versus Beta 1 comparison. Dashed line indicates threshold used [log_2_ (fold change) = 0.25].

### Oleate preferentially directs **β** cells towards proliferation while amplifying oxidative phosphorylation and antioxidant pathways and diminishing **β**-cell maturation

Growing evidence suggests that β cells are less differentiated during cell cycle activation (Sachs et al., 2020). Thus, we used CytoTRACE, a computational framework based on the number of transcripts expressed per cell (Gulati et al., 2020), to predict the differentiation state of cells from our scRNA-seq data set. We observed that the Beta 4 and 5 clusters were more differentiated, while the proliferative Beta 2 (including part of Beta 1 and 3 clusters) were less differentiated (Figure 5A). Interestingly, both palmitate and oleate treatment increased the immaturity score relative to vehicle (Figure 5B), suggesting that both FA promote dedifferentiation of β cells in agreement with the DEG analyses shown in Figure 2.

**Figure 5.**
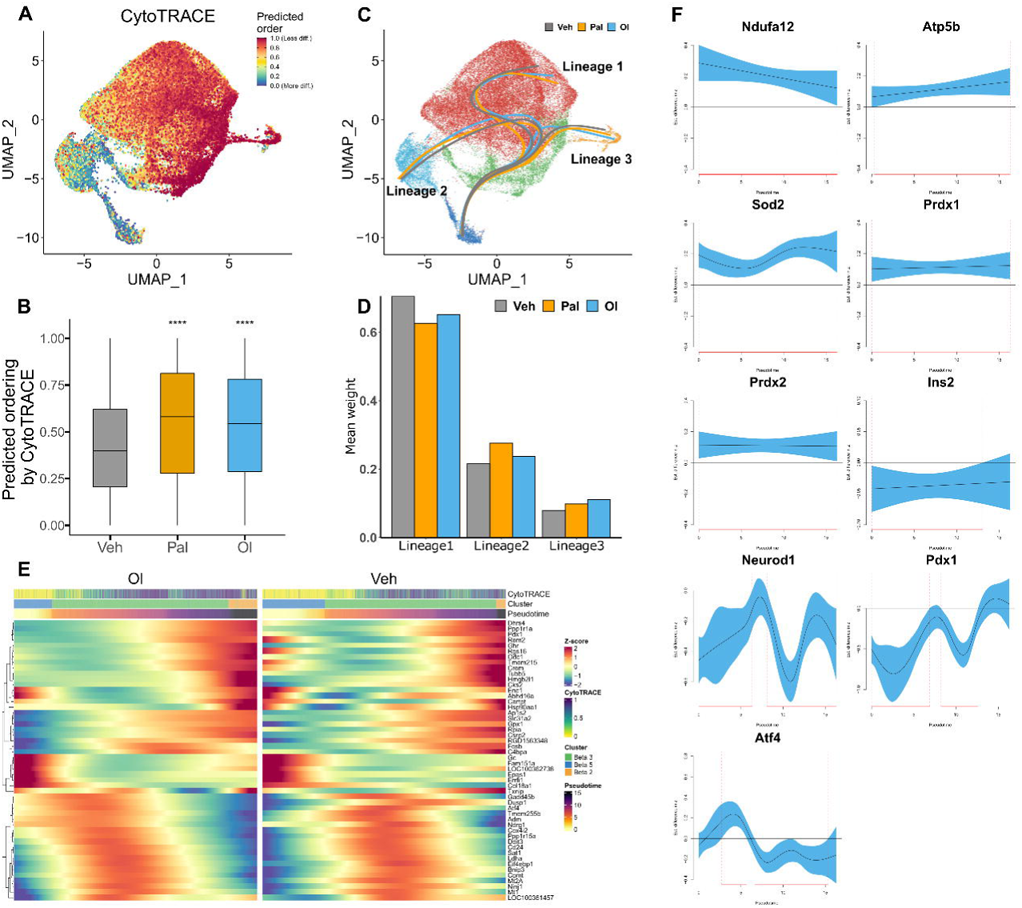
Trajectory inference analysis of β cells. **(A)** UMAP plot of pooled (Veh, Pal and Ol conditions) β cells showing the CytoTRACE score from the most (blue) to the least (red) differentiated cells. **(B)** Quantification of CytoTRACE predicted score for each of Veh, Pal and Ol conditions. Boxplot shows median, quartile and whisker values. **** indicates p<0.001 following one-way ANOVA with Dunnett’s multiple comparisons test. **(C)** UMAP plot showing the different pseudotime trajectories for each of Veh, Pal and Ol condition. **(D)** Average weight of each lineage for each of Veh, Pal and Ol conditions.**(E)** Heatmap showing the expression of the 50 most modulated genes along Lineage 3 in the Ol versus Veh comparison. **(F)** Generalized additive model plots of selected genes along Lineage 3 showing the expression level in the Ol condition (blue) relative to Veh (Y=0). Significant differences in expression levels (red line) are indicated along the pseudotime.

We then asked whether a transition state could be identified between the more differentiated Beta 5 cluster and the less differentiated Beta 2 cluster. To test this, we performed trajectory inference analysis across our multiple conditions using the Slingshot and Condiments R/Bioconductor packages (Street et al., 2018, De Bézieux et al., 2021), with a root starting at cluster Beta 5. Pseudotime ordering of β cells from the most differentiated cluster revealed 3 lineages (Figure 5C). These trajectories branched within the UPR/ER-stress-related Beta 3 subpopulation, leading to the secretory Beta 1 (lineage 1), the more differentiated insulin-expressing Beta 4 (lineage 2), or the proliferative Beta 2 (lineage 3) subpopulations. Although no lineage was unique to a particular condition, comparing the mean cell weight of each lineage calculated using Condiments (De Bézieux et al., 2021) across the 3 conditions showed that oleate favored the progression towards the proliferative path (lineage 3 has significant greater weights for oleate); that vehicle-treated islets differentiated preferentially through lineage 1; and that palmitate-treated islet cells differentiated preferentially through lineage 2 (Figure 5D). These results confirm that oleate conducts more cells towards proliferation.

Next, we identify genes whose expression patterns along the path to proliferation (lineage 3) are differentially modulated between the culture conditions. For each gene and for each experimental condition, smoothers were fitted using a negative binomial generalized additive model. Significant differences between smoothers were identified by comparing smooths in a factor-smooth interactions model. Compared to vehicle, expression of several genes was significantly modulated in the palmitate and oleate groups (Suppl. Table 7) and were involved in UPR/ER-stress related pathway (Suppl. Table 8).

As visually interpreting the differential modulation of gene expression along a pseudotime lineage between conditions is challenging with heatmaps (Figure 5E, Suppl. Figure 3A), the differences between pairs of smoothers were visualized in Figure 5 and Supplementary Figure 3B. In lineage 3, genes involved in the electron transport chain, such as *Ndufa12* and *Atp5b,* as well as the antioxidants *Sod2*, *Prdx1* and *Prdx2,* were upregulated, whereas β cell markers, such as *Ins2*, *Neurod1* and *Pdx1* were reduced in response to both oleate and palmitate compared to vehicle (Figure 5F, Suppl. Figure 3B). *Atf4*, associated with UPR/ER-stress, was up-regulated at the beginning of the pseudotime while down-regulated at the end, in both FA-exposed groups (Figure 5F, Suppl. Figure 3B).

### ROS generation, peroxiredoxin activity and MYC are necessary for oleate-induced β-cell proliferation

The potential role of ROS and antioxidant signaling in β cell proliferation in response to oleate suggested by scRNA-seq was experimentally tested in isolated rat islets. We focused on the peroxiredoxin family since *Prdx1, Prdx2, Prdx3* and *Prdx4* were significantly upregulated in the proliferative Beta 2 cluster as well as along the pathway to proliferation (lineage 3) in response to oleate (Suppl. Table 2, Figures 4 & 5). First, we measured β-cell proliferation in rat islets exposed oleate with or without the antioxidant N-acetylcysteine (NAC). NAC treatment dose-dependently reduced β cell proliferation in response to glucose + oleate but not glucose alone (Figure 6A). Then we exposed islets to Conoidin A, a specific inhibitor of peroxiredoxins (Stancill et al., 2020). Addition of Conoidin A dose-dependently decreased β-cell proliferation in response to glucose + oleate, but not glucose alone (Figure 6B). To confirm the selectivity of Conoidin A we tested whether it altered the proliferative response to the DYRK1A inhibitor harmine (Dirice et al., 2016). At the maximal concentration used in Figure 6B, Conoidin A reduced the response to glucose + oleate but not to harmine (Figure 6C).

**Figure 6.**
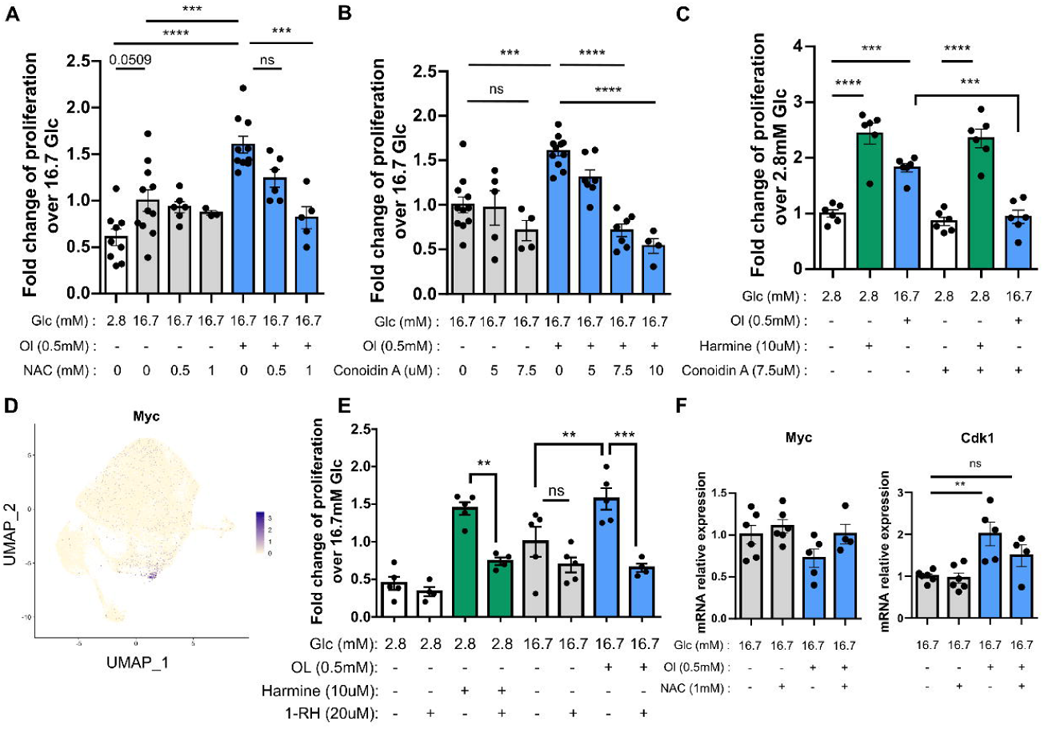
Functional analysis of ROS, peroxiredoxin and MYC in Ol-induced β-cell proliferation. **(A-C)** Isolated rat islets were exposed to 2.8 or 16.7 mM glucose, oleate (Ol; 0.5 mM) or harmine (10 μM) with or without NAC (0.5 or 1 mM), Conoidin A (5 or 7.5 μM) or vehicle as indicated. Beta-cell proliferation was assessed by flow cytometry following staining for EdU and C-peptide and presented as the fold-change of the percentage of EdU^+^/C-peptide^+^ over total C-peptide^+^ over the control (16.7 mM glucose) condition. **(D)** UMAP plot of pooled (Veh, Ol and Pal conditions) β cells showing the expression of *Myc*. **(E)** Isolated rat islets were exposed to 2.8- or 16.7-mM glucose, oleate (Ol; 0.5 mM) or harmine (10 mM) with or without 1-RH (20 μM) or vehicle as indicated. β-cell proliferation was assessed as above. **(F)** Isolated rat islets were exposed to 16.7 mM glucose with or without oleate (Ol; 0.5 mM), NAC (1 mM) or vehicle as indicated. *Myc* and *Cdk1* mRNA were quantified by RT-PCR and normalized to cyclophilin. Data are presented as the fold-change over the control condition (16.7 mM glucose). Data represent individual values and are expressed as means +/− SEM. **p<0.01, ***p<0.005, ****p<0.001 following one-way ANOVA with Tukey’s (A, B, E), Sidak’s (C) or Dunnett’s **(E)** multiple comparisons test. NS, not significant.

The proto-oncogene MYC plays a key role in β-cell proliferation (Rosselot et al., 2021, Rosselot et al., 2019) and is a potential effector of ROS signaling (Elouil et al., 2005, Benassi et al., 2006). We therefore asked whether MYC was necessary for β-cell proliferation in response to oleate. Intriguingly, we identified a small group of *Myc*-expressing cells within the Beta 3 subpopulation (Figure 6D) and MYC target genes (e.g. *Cdk1*, *Cdc20*, *Ccna2*) were upregulated in proliferating β cells (Suppl. Table 2). In rat islets exposed to the small molecule inhibitor of the MYC/MAX interaction 1-RH (Zhang et al., 2010) we observed a decrease in the β cell proliferative response to both harmine and glucose + oleate (Figure 6E). Accordingly, oleate increased expression of the MYC target gene *Cdk1*, but not *Myc*, and NAC treatment blocked this effect (Figure 6F). Together, these results show that ROS generation and peroxiredoxin activity are necessary for oleate-induced β-cell proliferation and that ROS likely acts via MYC.

## DISCUSSION

The aims of this study were to ascertain the differential effects of and transcriptional response to individual FA on β-cell proliferation, and to characterize the transcriptome of proliferating β-cells in response to FA at single-cell resolution. Palmitate and oleate were selected a representative unsaturated and monounsaturated species, respectively, because they are the 2 most abundant FA in human plasma (Abdelmagid et al., 2015). scRNA-seq enabled us to capture several β ell clusters, including proliferating cells. Both FA increased energy metabolism and decreased expression of β-cell maturation markers. Notable differences between the two FA included a marked increase in cellular stress in response to palmitate, but not oleate. Comparing the transcriptomic profile of proliferative and non-proliferative cells at the gene and trajectory levels highlighted a key role for ROS and peroxiredoxin signaling in the path to proliferation. Accordingly, ROS scavenging or peroxiredoxin inhibition blocked oleate-induced β cell proliferation, a process that requires MYC activity.

β-cell functional heterogeneity is well established, and single-cell transcriptional profiling studies have reinforced this notion by identifying multiple β-cell subtypes with unique transcriptional signatures [reviewed in (Wang and Kaestner, 2019, Mawla and Huising, 2019)]. In this study, independent of the culture condition, we captured 5 transcriptionally distinct β cell clusters characterized by high expression of maturity and/or function (Beta 1, 4 and 5), proliferative (Beta 2) and UPR/ER stress (Beta 3) markers. Although β cell clusters distinguished by expression of ER and oxidative stress as well as maturity markers were identified previously in human β cells (Baron et al., 2016, Muraro et al., 2016), not surprisingly transcriptional signatures and β-cell clustering differed significantly in our study likely due to species, culture and methodological differences but also to the limitations of the scRNA-seq technology (Mawla and Huising, 2019).

Differential effects of individual FA on the β cell have been investigated in the context of chronic exposure to elevated levels of glucose and FA, so called glucolipotoxicity, which leads to β-cell dysfunction, apoptosis and dedifferentiation through various mechanisms including ER and oxidative stress, inflammation, and impaired autophagic flux [reviewed in (Lytrivi et al., 2020a)]. In this context, saturated FA (e.g. palmitate) have deleterious effects on β-cell function and survival (El-Assaad et al., 2003), while unsaturated FA (e.g. oleate) are protective (Sargsyan et al., 2016, Maedler et al., 2001, Maedler et al., 2003). The advent of unbiased transcriptomic technologies has enabled examination of the effects of FA on islet function at the level of global changes in gene expression. Microarray or bulk RNA-seq approaches of human islets as well as rodent insulin-secreting cell lines have shown that exposure to palmitate alone or in the presence of high glucose concentrations leads to a gene expression profile similar to what is observed in islets from type 2 diabetic donors, characterized by an increase in UPR/ER-stress, protein degradation, immune surface receptor, autophagy, mRNA splicing regulation, and nuclear transport processes (Lytrivi et al., 2020b, Bugliani et al., 2013, Cnop et al., 2014, Hall et al., 2014, Marselli et al., 2020). To our knowledge, our study is the first to compare transcriptomic profiles of islets exposed to individual FA at the single-cell level. Thus, in line with previous transcriptomic analyses of whole islets, we found that genes involved in UPR/ER-stress were upregulated in the Beta 3-5 subpopulations in response to palmitate, as well as translational processes and oxidative stress (Carlsson et al., 1999, Barlow and Affourtit, 2013, Hatanaka et al., 2014), confirming the deleterious effects of saturated FA. Interestingly, not all β-cell subpopulations were affected (e.g. Beta 1 and 2); however, whether heterogeneity described here is a snapshot of a dynamic flux or a stable cell state remains to be explored. On the other hand, cellular stress responses were not augmented in response to oleate, in agreement with previous studies (Maedler et al., 2001, Maedler et al., 2003, Sargsyan et al., 2016). Instead, both oleate and palmitate increased expression of genes involved in glycolysis and oxidative phosphorylation in several β-cell clusters (Beta 1, Beta 4, and Beta 5). Intriguingly, these results resemble the transcriptional response of the human β cell in diabetes (Camunas-Soler et al., 2020) and may reflect an attempt by the β cell, albeit failed in the case of palmitate, to meet the elevated secretory demand.

Although no β−cell cluster was unique to an individual culture condition, the relative proportion of each cluster was dependent on the specific FA treatment. We observed a higher proportion of proliferative β cells in oleate-treated islets. Furthermore, we report distinct maturity states across the different clusters, especially in the path toward proliferation, where proliferative β cells were less mature than the others. These data are consistent with other studies where cycling β cells show immature gene expression profiles (Sachs et al., 2020, Bader et al., 2016). Interestingly, both oleate and palmitate decreased expression of β-cell markers and the overall maturity score. Although the effect of palmitate is in line with previous studies (Hagman et al., 2005, Busch et al., 2002, Kim-Muller et al., 2016), β-cell dedifferentiation following chronic exposure to glucose and oleate has not, to our knowledge, been described previously.

By comparing the transcriptome of proliferative (Beta 2) to non-proliferative β cells we found that besides the expected upregulation of pathways involved in cell division processes, proliferative β cells showed upregulated genes and pathways related to oxidative phosphorylation and antioxidant activity. Importantly, ROS and peroxiredoxins were found to be essential for oleate-induced β-cell proliferation. Nutrients are important inducers of ROS (e.g., O_2_ and H_2_O_2_) (Roma and Jonas, 2020) in the β cell, and although excessive ROS production is associated with dysfunction (Tanaka et al., 1999, Tanaka et al., 2002), a moderate increase in ROS levels is required for glucose-induced insulin secretion (Pi et al., 2007), promotes β-cell neogenesis in zebrafish (Ahmed Alfar et al., 2017) and postnatal β-cell proliferation in mice (Zeng et al., 2017). Hence, the increase in peroxiredoxin levels, documented in this study, may be important to limit ROS, but also to function as signal mediators by oxidizing target thiols in a redox relay (Roma and Jonas, 2020).

In response to nutrient excess, ROS are generated in the β-cell mitochondria (Carlsson et al., 1999, Escribano-Lopez et al., 2019), peroxisome (Elsner et al., 2011) and ER (Melo et al., 2017). Although the increased expression of genes associated with oxidative phosphorylation suggests a mitochondrial origin, other sites of ROS generation may contribute to β-cell proliferation. Among other possibilities, sustained insulin demand induces a β cell proliferative response through moderate activation of the UPR (Sharma et al., 2015), suggesting that ROS may also be produced in the ER during insulin processing. Supporting this hypothesis, pseudotime ordering of cells suggested that the path towards proliferation (Beta 2) goes through the UPR/ER-stress associated Beta 3 population. Furthermore, genes significantly modulated by FA along this path include UPR/ER-stress-related genes such as *Atf4*, known to also trigger an antioxidant response (Harding et al., 2003). Further studies will be required to establish the subcellular origin of ROS and the link to peroxiredoxins in response to oleate.

Consistent with the important role of MYC in β-cell proliferation (Rosselot et al., 2019), we found that MYC inhibition decreased β-cell proliferation in response to oleate. *Myc* was expressed in a small group of cells in the path towards proliferation and MYC target genes (e.g., *Cdk1*, *Cdc20*, *Ccna2*) were upregulated in proliferating β cells. In agreement with these findings, nutrients increase *Myc* expression in islets in vivo and in vitro (Jonas et al., 2001). Upstream of MYC, ROS promotes *Myc* expression in islets (Elouil et al., 2005) and ERK-dependent phosphorylation in melanoma cells (Benassi et al., 2006). We showed that ROS signaling is also important for MYC target gene expression in response to oleate, revealing a potential mechanistic link between oleate-induced ROS and the β-cell proliferative response.

In summary, our findings suggest that ROS and peroxiredoxin signaling via MYC is required for oleate-induced β-cell proliferation in rat islets, providing important information on the molecular mechanisms of β-cell adaptation to nutrient excess.

## ACKNOWLEDGEMENTS

We thank Grace Fergusson and Mélanie Éthier from the Rodent Metabolic Phenotyping core of the CRCHUM for assistance with islet isolations; Dominique Gauchat and Philippe St-Onge from the Flow Cytometry core of the CRCHUM for assistance with measurements of β cell proliferation. This study was supported by the National Institutes of Health (R01-DK-58096), the Canadian Institutes of Health Research (MOP 77686), and the Quebec Cardiometabolic Health, Diabetes and Obesity Research Network.

## AUTHOR CONTRIBUTION

A.V: Conceptualization, supervision, methodology, investigation, formal analysis and writing - original draft. J.G: Conceptualization, methodology, investigation, writing, editing. A.FM: Methodology, investigation, formal analysis, writing. ZE.A: Investigation, formal analysis. AL.C: Methodology. R.S: Conceptualization, validation. V.P: Conceptualization, validation, writing, editing, funding acquisition, and project administration. V.P is the guarantor of this work and, as such, had full access to all the data in the study and takes responsibility for the integrity of the data and the accuracy of the data analysis.

## METHODS

### Reagents and Solutions

RPMI-1640, HBSS, and FBS were purchased from Life Technologies Inc. (Burlington, ON, Canada). Penicillin/Streptomycin and D-PBS were from Multicell Wisent Inc. (Saint-Jean-Baptiste, QC, Canada). FA-free BSA was from Equitech-Bio (Kerrville, TX, USA). Histopaque-1119/1077, glucose, harmine, sodium oleate and palmitate were from Sigma Aldrich (Saint-Louis, MI, USA). NAC and Conoidin A were from Cayman Chemical (Ann Arbor, MI, USA). 10058-F4/1-RH was from Tocris (Bristol, United Kingdom).

### Animals

All procedures were approved by the Institutional Committee for the Protection of Animals at the Centre Hospitalier de l’Université de Montréal. Male Wistar rats weighing 250–300 g (~2 months old) (Charles River, Saint-Constant, QC, Canada) were group housed (2 animals per cages) under controlled temperature on a 12 h light-dark cycle with free access to water and standard laboratory chow.

### Islet isolation

Islet isolation was performed by collagenase type XI (Sigma-Aldrich, Oakville, ON, Canada) digestion of the pancreas as described (Jacqueminet et al., 2000). In brief, 10 ml of collagenase [0.65mg/ml in Hanks buffered saline solution (HBSS)] was injected into the bile duct. The perfused pancreas was dissected and placed in water bath for 10-13 min at 37 °C. Then, 25 ml of cold HBSS+BSA (1%) was added to the samples prior to mechanical digestion by hand shaking, followed by centrifugation at 339xg at 4°C. Samples were then mesh screen filtered and washed twice with HBSS+BSA solution. Pellets were resuspended in 10 ml of Histopaque-1119. Second (Histopaque-1077) and third layer (HBSS+BSA) were respectively added to form a 3-layer gradient. Samples were centrifuged at 339xg for 30 min at 4°C. The interphase between the upper and the middle layers of the gradient was harvested and washed once with HBSS+BSA. Islets were hand-picked under the microscope prior to culture.

### Islet culture

Following isolation islets were allowed to recover for 24 h in RPMI-1640 with 10% (vol./vol.) FBS (Invitrogen, Burlington, VT, USA) with 11.1 mM glucose. For treatment, pools of 200 islets were cultured in RPMI-1640 with FBS for 48 h in the presence of glucose (2.8 mM or 16.7 mM) with or without palmitate (palmitate) or oleate (oleate) (0.5 mM) or the corresponding vehicle (Veh) [0.1 mmol / l BSA + 50% ethanol (vol./vol.)] as indicated in the figure legends. FA were dissolved in ethanol prior to 1-h complexation with BSA at a 5:1 ratio (FA:BSA). To investigate cell proliferation, 5-Ethynyl-2’-deoxyuridine (EdU; 10 µM) was added to the culture medium. Media were replaced daily.

### Proliferation by flow cytometry

Islets were treated for 48 h as described above with or without NAC, Conoidin A, 10058-F4/1-RH or harmine as indicated in the figure legends. After treatment, islets were washed once with D-PBS + 2 mM EDTA solution (339xg, 3 min, 4°C) and digested by enzymatic disaggregation for 10 min at 37°C using Accutase (Innovative Cell Technologies Inc., San Diego, CA, USA). The reaction was stopped using islet culture medium and the islet cell suspension was washed once with PBS and dead cells labeled using the LIVE/DEAD™ Fixable Aqua Dead Cell Stain Kit (405 nm; Invitrogen Inc., USA). Sample fixation, permeabilization and EdU detection were performed using the Click-iT™ Plus EdU Alexa Fluor™ 488 Flow Cytometry Assay Kit (BD Biosciences, San Jose, CA) according to the manufacturer’s instructions. Labeling of β cells was performed by incubating samples for 30 min with anti-C-peptide and anti-glucagon antibodies, respectively. Fluorophore-coupled primary antibodies and dilutions are listed in Supplementary Table 1. Dead-cell stain, EdU, C-peptide and Glucagon labelled cells were detected on an LSRII flow cytometer with FACSDiva™ Software (BD Biosciences, San Jose, CA) using the 405-, 488-, 640- and 561-nm lasers coupled with 525/50-, 530/30-, 670/14- and 586/15-nm BP filters. Proliferation was calculated as the percentage of double-positive EdU^+^ and C-peptide^+^ cells over the total C-peptide^+^ cell population. A minimum of 10,000 C-peptide^+^ cells were counted in each sample. Results were presented using FlowJo v10.7 software (Ashland, OR; https://www.flowjo.com/solutions/flowjo).

### Quantitative PCR

RNA was extracted from 150-200 islets using the RNeasy Micro kit (Qiagen, Valencia, CA). RNA was quantified by spectrophotometry with a NanoDrop 2000 (Life Technologies Inc.), and 1 µg of total RNA was reverse transcribed using M-MLV reverse transcriptase (Thermo Fisher Scientific, Waltham, MA). Real-time PCR was performed using the QuantiNova SYBR Green RT-PCR Kit (Qiagen, Valencia, CA). Results are expressed as the ratio of target mRNA to cyclophilin-A RNA levels and normalized to the levels in control islets. Primers sequences are listed in Supplementary Table 1.

### Statistical analysis (excluding scRNA-seq data)

All statistical analyses were performed using GraphPad Prism 9. We used the one-way analysis of variances (ANOVA) followed by Tukey’s, Dunnett’s or Sidak’s post hoc analysis for multiple comparisons to determine significance among the different treatment groups as indicated in the figure legends. Results are presented as mean ± SEM. p < 0.05 was considered significant.

### scRNA-sequencing

Prior to scRNA-seq library generation, islets were dissociated (see above), washed once using PBS+BSA (1%) solution and resuspended with MACS MicroBeads (Miltenyi Biotec, Waltham, MA). Dead cells were removed using MS Columns (Miltenyi Biotec) according to the manufacturer’s instruction. Live cells were then counted using a Countess II automated cell counter (Thermo Fisher Scientific, Waltham, MA). Single cell suspensions with <10% dead cells were kept for library generation. Single-cell libraries were generated using the Chromium Single-cell 3′ library and gel bead kit v2 (10x Genomics, Pleasanton, CA). In brief, to reach a target cell number of 6,000 cells per sample, 10,500 cells per sample were loaded onto a channel of the 10x chip to produce Gel Bead-in-Emulsions (GEMs). This underwent reverse transcription to barcode RNA before cleanup and cDNA amplification followed by enzymatic fragmentation and 5′ adaptor and sample index attachment. Library quantification and quality control was performed on a 2100 bioanalyzer (Agilent, Santa Clara, CA). Libraries were sequenced on the NovaSeq 6000 (Illumina, San Diego, CA) with 150 bp paired-end sequencing.

### Preprocessing of droplet-based scRNA-seq data

Demultiplexing of binary base call (BCL) files, alignment to the *Rattus norvegicus* genome (version Rnor_6.0), read filtering, barcode and unique molecular identifier (UMI) counting were performed using the CellRanger analysis pipeline (v2.0) provided by 10x Genomics. High quality barcodes were selected on the basis of the overall distribution of total UMI counts per cell using the standard CellRanger cell detection algorithm. All further analyses were run with R (version 4.1.0) using the Seurat R package (v4.0, https://satijalab.org/seurat/). We excluded genes expressed in less than 4 cells from the analysis; as well as cells that had a high fraction of counts from mitochondrial genes (20% or more), that expressed fewer than 200 genes, or that had more than 4,000 genes. Based on these criteria, 2,241 cells (from 54,811 cells) were discarded, resulting in 52,570 cells for further analysis. We did not see bias in the number of cells excluded based in the culture condition.

### Embedding, clustering and cell type annotation

Sample Raw counts were normalized separately using SCTransform (Hafemeister and Satija, 2019) and integrated using Seurat (Stuart et al., 2019). The normalized and log-transformed expression values were used for downstream analysis. PCA was performed on the normalized expression using top 3,000 most variable genes across all samples. A neighborhood graph was built with n-neighbors set to 20 and 30 calculated PCs as inputs. This neighborhood graph was used as input for clustering via the Leiden algorithm (with a resolution parameter of 0.4) (Traag et al., 2019) and the identified clusters were visualized using Uniform Manifold Approximation and Projection (UMAP) (McInnes et al., 2020). Annotation of cell types was performed using a manually curated list of previously characterized cell type markers (e.g., *Ins2, Gcg, Sst and Ppy* for Beta, Alpha, Delta, and PP-cells respectively). Re-clustering of the Beta Cell subpopulation (39,910 cells), identified in the first round of clustering, was performed by extracting the matrix of raw counts of transcripts followed by a second round of normalization, sample dataset integration, dimension reduction and clustering via the Leiden algorithm (with a resolution parameter of 0.4).

### Differentially expressed genes and pathway analysis

Pairwise differential gene expression comparisons were made across experimental treatment groups and/or cell type subpopulation. Scaled normalized expression values were used as input to Seurat’s FindMarkers function implementing the MAST test (Finak et al., 2015) to identify DEGs. Genes were declared significantly differentially expressed at a False Discovery Rate (FDR) cutoff of 5% and absolute log2 fold change > log2(1.2). MsigDB’s Hallmark and C5 (Gene Ontology) gene sets were used for Fisher Enrichment Analysis followed by an Enrichment Map (Merico et al., 2010) analysis to aggregate and organize overlapping GO gene-sets into networks making functional interpretation easier. Gene sets were declared significant at an FDR of 5%. The CytoTRACE function from the R package CytoTRACE (Gulati et al., 2020) was used to assign a differentiation state score to each cell based on expression of a set of genes.

### Trajectory inference

The Slingshot (Street et al., 2018) R/Bioconductor package was used to reconstruct lineages and infer pseudotime. The goal of Slingshot is to use clusters of cells to uncover global structure and convert this structure into smooth lineages represented by a smooth curve and a one-dimensional variable, called pseudotime. A single lineage represents a path in a cellular trajectory (from a start point to end point) and the cells belonging to the lineage are then ordered by the pseudotime calculated by Slingshot. The Condiments R/Bioconductor package (De Bézieux et al., 2021), which implements a statistical workflow for modeling cell trajectories across multiple experimental conditions was used to refit a slingshot trajectory for each experimental condition and to map the Cell weights, the probability that a cell belongs to a particular lineage in the trajectory, to each experimental condition.

To identify genes whose expression patterns along the trajectory are differentially modulated between the experiment’s conditions, for every gene in the dataset we fit a negative binomial generalized additive model (NB-GAM), using the gam function as implemented in the R’s package mgcv (doi: 10.1201/9781315370279), to smooth each gene’s expression in each lineage and generate separate smoothers for each condition based on the pseudotime inferred by slingshot. We tested whether the smoothers are significantly different between conditions by comparing smooths in factor-smooth interactions. The resulting p-values were adjusted for multiple testing and significance set at FDR level set to 5%. Significant Differences between pairs of smooths were visualized using plot_diff function as implemented in R’s package itsadug.

## RESOURCE AVAILABILITY

### Lead contact

Further information and requests for resources and reagents should be directed to and will be fulfilled by the lead contact, Vincent Poitout

### Materials availability

This study did not generate new unique reagents.

### Data and Code Availability

scRNA-sequencing data will be submitted to the NCBI Gene Expression Omnibus (GEO; https://www.ncbi.nlm.nih.gov/geo/) and original code will be available as supplemental data file prior to publication.

## SUPPLEMENTAL INFORMATION

Supplementary document 1: Suppl. Table 1 and Suppl. Figures 1–3

Supplementary Table 2

Supplementary Table 3

Supplementary Table 4

Supplementary Table 5

Supplementary Table 6

Supplementary Table 7

Supplementary Table 8

