## Supplementary document 1 for "Single-cell RNA sequencing reveals a role for reactive oxygen species and peroxiredoxins in fatty-acid-induced rat ß-cell proliferation"

**Supplementary Table 1**

**Flow cytometry antibodies**

| Antibody | Dilution/Concentration | Company | Cat # |
|---|---|---|---|
| Alexa Fluor® Mouse anti-C-peptide | 1:25 | BD Biosciences | 565831 |

**RT-PCR primers**

| | Forward sequence | Reverse sequence |
|---|---|---|
| Cyclophilin-A | GCCATTATGGCGTGTGAAGTC | CTTGCTGCAGACATGGTCAAC |
| Myc | GAGGTGGAAAACCCGACAGT | AAATAGGGCTGCACCGAGTC |
| Cdk1 | GGAACAGAGAGGGTCCGTTG | CCACACCATAAGTCCCTTCTCC |

### Supplementary Figure 1

A

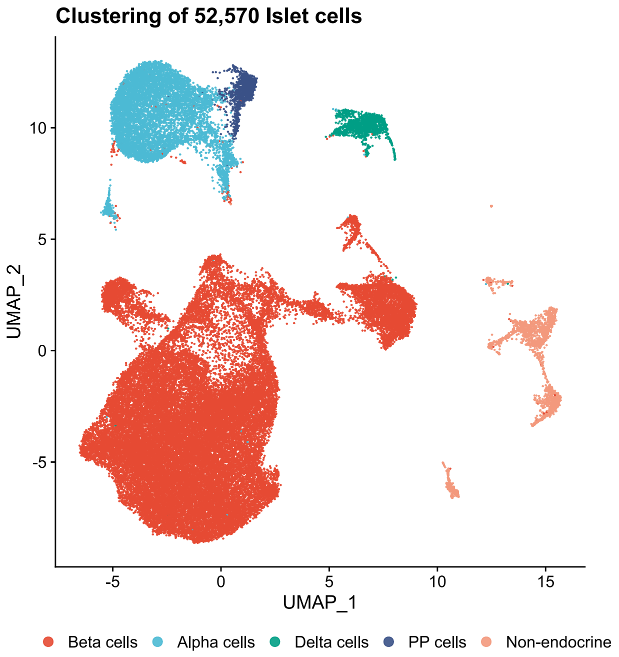


B

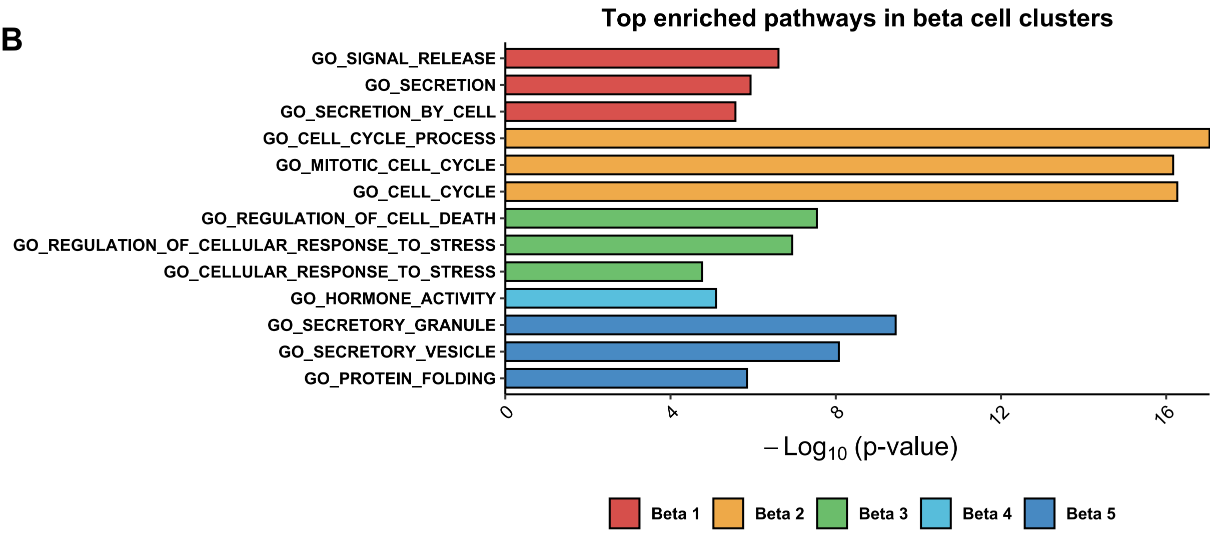


**Supplementary Figure 1. Single cell transcriptome analyses of rat islets and pathway enrichment analysis of β-cell subpopulations.**
(A) UMAP plot of pooled (Veh, Pal and Ol conditions) islet cells (52,570 cells in total) indicating endocrine (α, β, δ and PP cells) and non-endocrine cell types.
(B) Selected Gene Ontology (GO) terms from pathway enrichment analyses from the top 50 β-cell subpopulation markers (pathways listed in Suppl. Table 3).

#### Supplementary Figure 2

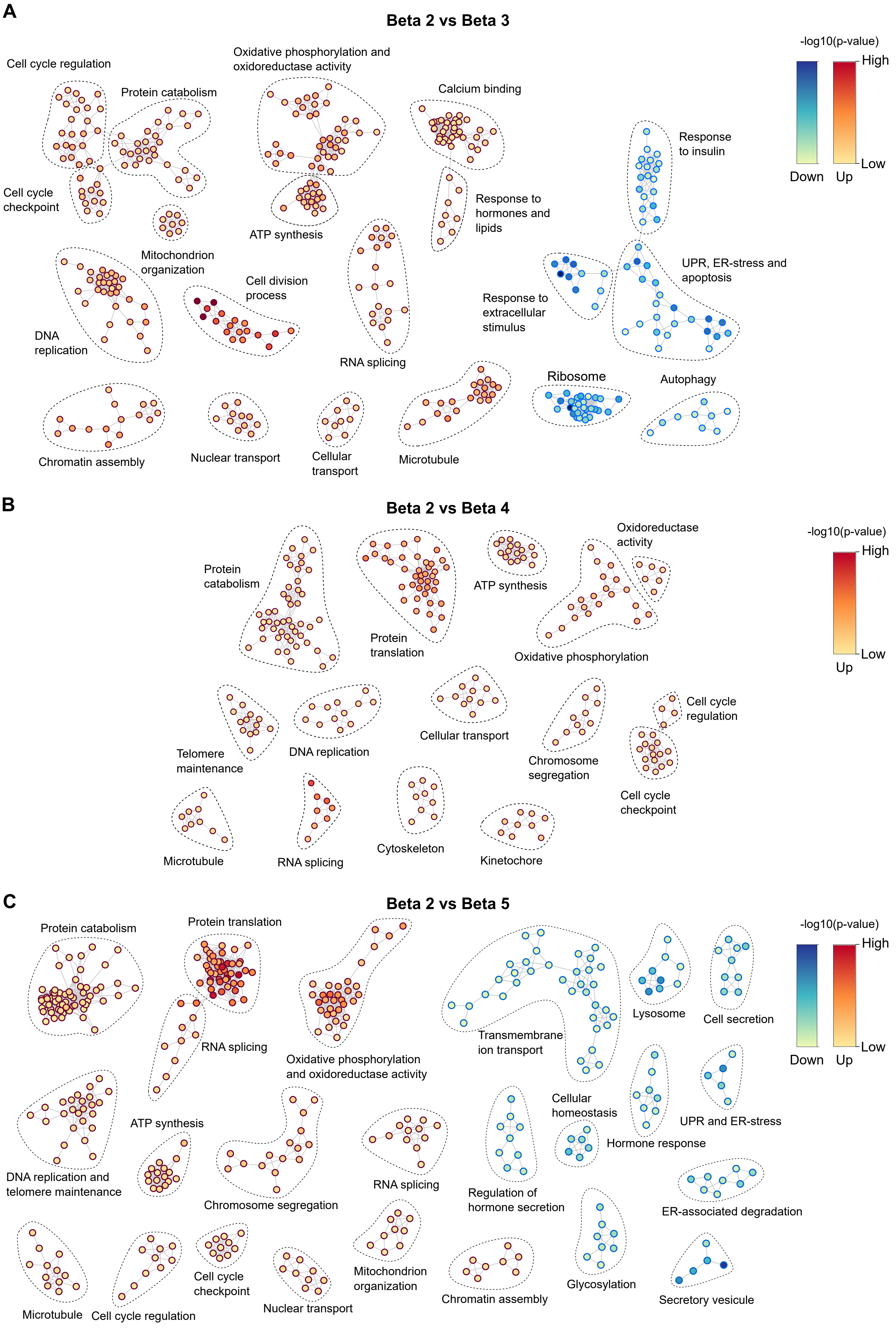


**Supplementary Figure 2. Comparison of proliferative to non-proliferative β cells.** (A-C) Pathway enrichment map of pooled (Veh, Pa and Ol conditions) proliferative Beta 2 versus non-proliferative Beta 3 (A), Beta 4 (B) and Beta 5 (C) subpopulations. Nodes represent pathways. Red or blue color gradient represents lower or higher significance for the up- and down-regulated pathways, respectively.

### Supplementary Figure 3

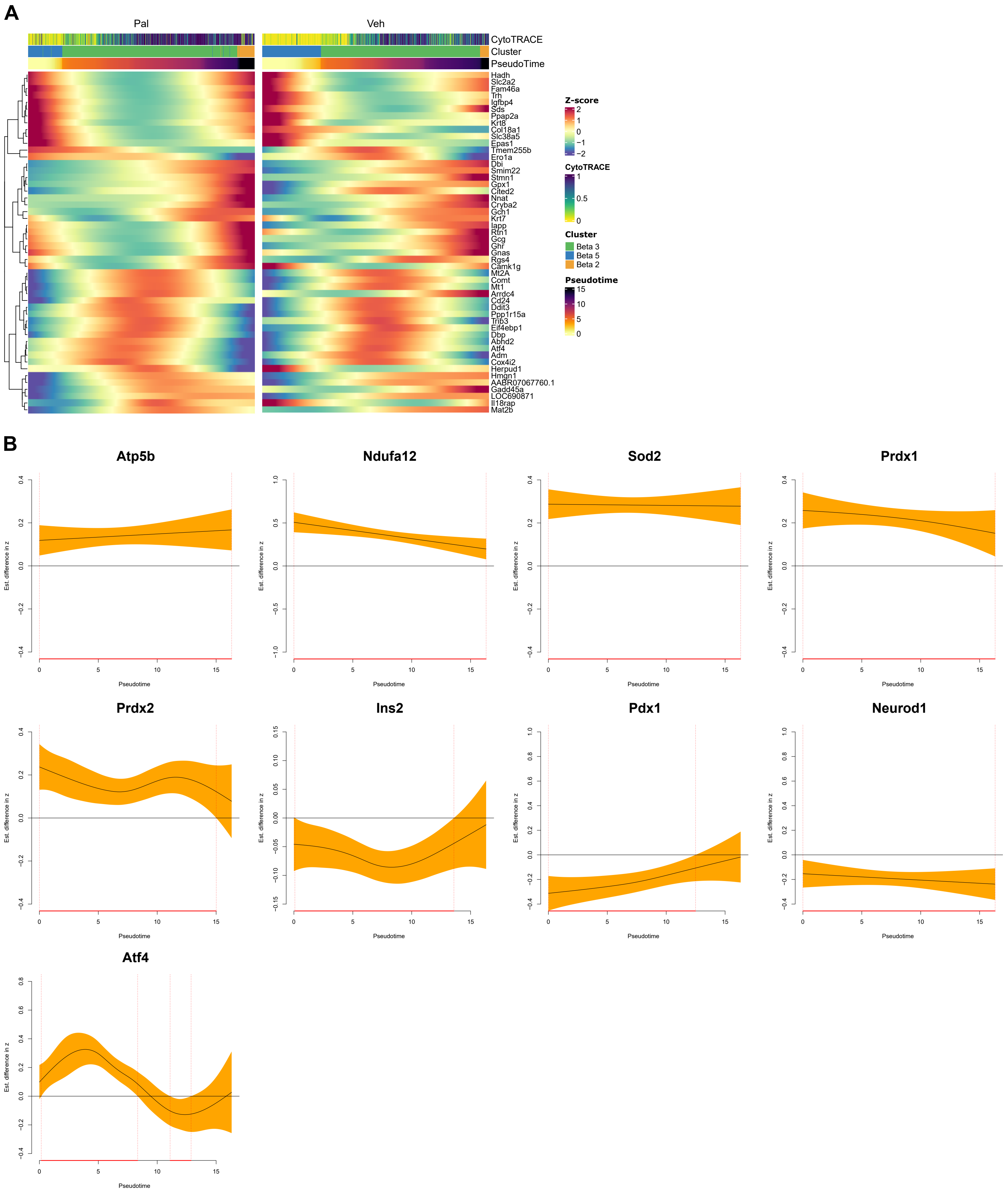


**Supplementary Figure 3. Trajectory inference analysis of β cells.**

(A) Heatmap showing the expression of the 50 most modulated genes along Lineage 3 in the Pal versus Veh comparison.

(B) GAM plots of selected genes along Lineage 3 showing the expression level in the Pal condition (yellow) relative to Veh (Y=0). Significant differences in expression levels (red line) are indicated along the pseudotime.
